# Rheology of crossbridge ensembles

**DOI:** 10.1101/2021.12.29.474482

**Authors:** Khoi D. Nguyen, Madhusudhan Venkadesan

## Abstract

How skeletal muscle responds to mechanical perturbations, its rheology, is important for animal movement control. The molecular machinery of myosin II-actin crossbridge cycling is a crucial part of muscle’s rheological properties, and multiple models have been proposed for this mechanochemical process. But current understanding of the scale-connection from individual molecular motors to ensemble rheology is limited. Here we present computational and mathematical analyses of several different hypotheses of crossbridge dynamics, from 2-state to 5-state myosin II motor models, and show that an ensemble of actomyosin crossbridges exhibits surprisingly simple rheological behavior in all cases. The ensemble rheology is captured by a sum of at most three linear viscoelastic sub-processes, and as few as one for some crossbridge models. This simplification lends itself to computationally efficient phenomenological muscle models with experimentally measurable parameters, while still remaining grounded in crossbridge theory. However, the collapse of the ensemble behavior to few linear sub-processes identifies major limitations of crossbridge models that cannot be resolved by adding complexity to the crossbridge cycle and point to the roles of inter-crossbridge interactions and non-crossbridge elements.

## Introduction

Muscle exerts force in response to *in vivo* neural inputs and length perturbations (Nishikawa et al., 2018; Zajac, 1989). Like most soft passive materials, the forces induced by the length perturbation, or simply perturbation response, relax over time (Huxley and Simmons, 1971; Nguyen et al., 2018) and the timescales involved are crucial for the functional role of muscle because they enable faster-than-reflex reactions (Bizzi et al., 1982; Hogan et al., 1987; Nishikawa et al., 2007; Nguyen et al., 2018, figure 1a). For example, on durations shorter than the relaxation timescales, the perturbation response is nearly unchanging and the muscle functions as an elastic solid body to resist stretching and aid in elastic energy storage in tendons (Bizzi et al., 1982; Biewener and Roberts, 2000). And on durations longer, the perturbation response relaxes significantly and the muscle instead functions like a viscous damper which enables rapid postural changes and kinetic energy dissipation (Lin and Rymer, 2000; Konow et al., 2012). But unlike passive materials, the muscle’s relaxation timescales are neurally regulated by active molecular interactions of myosin motors with actin filaments (Huxley and Tideswell, 1996), and whether a muscle functions like an elastic body, a viscous damper, or an viscoelastic intermediate is subject to neural control. Thus, understanding a muscle’s perturbation response in terms of its underlying molecular machinery helps to identify the interplay of neural inputs and viscoelastic rheology in an animal’s motor control of its muscles (Nguyen et al., 2018; Bizzi et al., 1982; Nishikawa et al., 2018, figure 1a).

**Figure 1:**
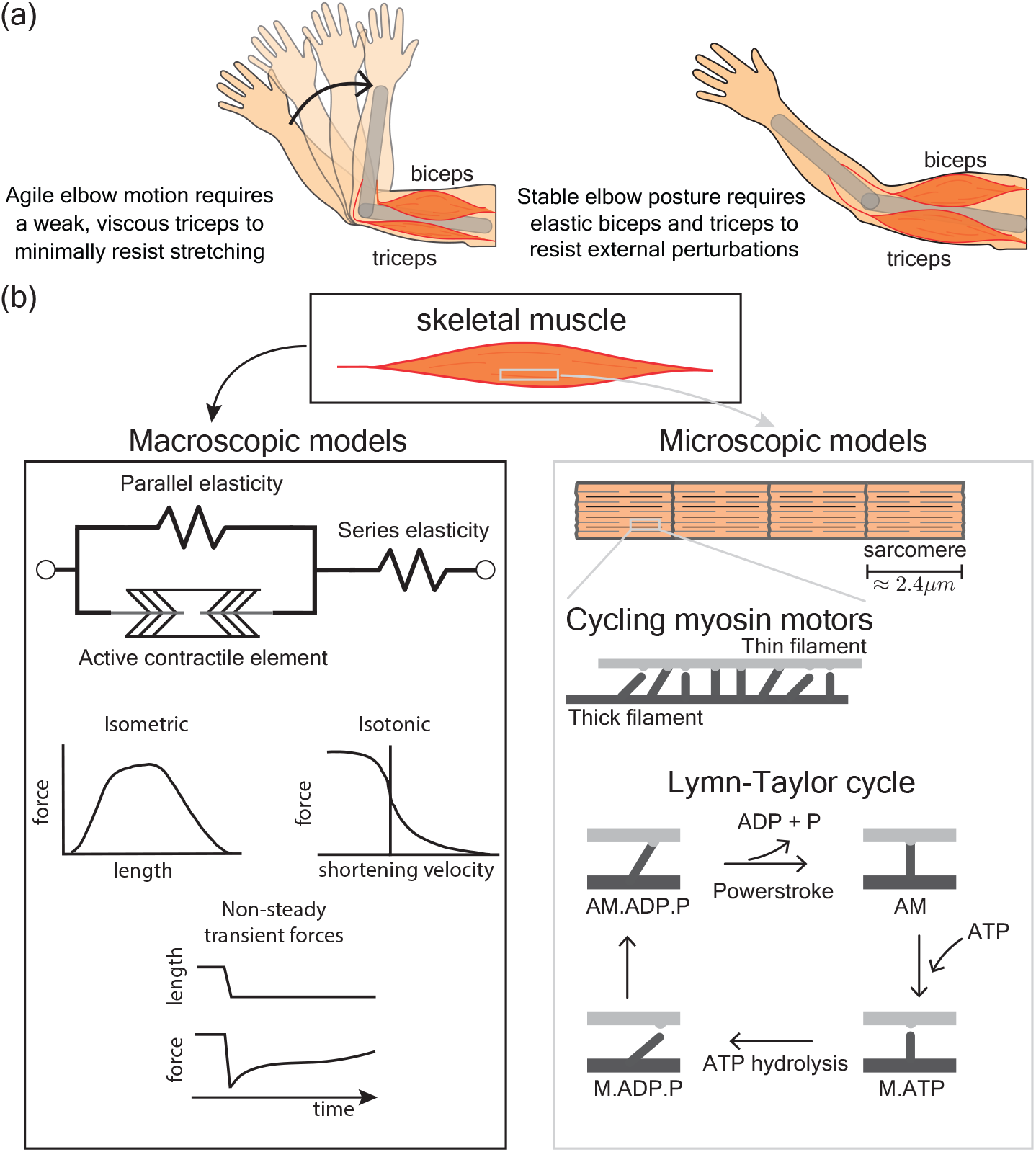
Current models of skeletal muscles. (a) An example of the motor functionalities provided by a muscle’s rheological response to perturbations. (b) Widely used muscle models can be separated into microscopic biochemical models and macroscopic phenomenological models. The microscopic models track the dynamics of myosin motors whereas macroscopic models apply isometric and isometric characterizations and transient forces to capture the active contractile element. The isometric and isotonic characterizations are adapted from figure 8 of Zajac 1989 and the transient forces are adapted from figure 3 of Huxley and Simmons 1971. The current paper seeks to connect the crossbridge cycle dynamics to the emergent generalized transient force response beyond the isometric and isotonic characterizations of muscle.

Muscle properties such as short-range stiffness, viscous damping, loss modulus, and storage modulus are examples of bulk rheological properties (Rack and Westbury, 1974; Konow et al., 2012; Kawai and Brandt, 1980) that are characterizations of the muscle’s perturbation response and are analogous to similar characterizations of passive materials (Tschoegl, 2012) and active materials (Gardel et al., 2004; Koenderink et al., 2009). The bulk rheological properties arise in part from the internal molecular machinery, similar to cytoskeletal networks (Grewe and Schwarz, 2020), and in part from muscle’s passive tissues and geometry (Sleboda and Roberts, 2020; Fung, 2013). The mapping from the microscopic dynamics of the molecular machinery to its contributions to bulk properties remains incomplete, however, and this gap is exemplified by the split between two classes of widely used muscle models. Namely, macroscopic phenomenological models and microscopic mechanochemical models that idealize muscles as one-dimensional structures.

Phenomenological models inspired by A.V. Hill’s work (Hill, 1938) are widely used to quantitatively describe muscle’s force production capabilities (Zajac, 1989; MacIntosh and MacNaughton, 2005; Fung, 1971; Todorov, 2003). These models incorporate an active contractile element, a parallel element that is often modeled as a passive elastic or viscoelastic material, and a series elasticity that captures tendon and other passive tissue compliance (figure 1b). The passive parallel element represents non-contractile tissues and the intramuscular fluid that have well-known effects on the contractile force capacity of muscle (Sleboda and Roberts, 2020). But muscle’s perturbation response is dominated by the active contractile element except under extreme eccentric conditions where the passive parallel element is important (MacIntosh and MacNaughton, 2005; Lindstedt et al., 2001). In Hill-type models, the perturbation response of the active element are represented by isometric force-length and isotonic force-velocity properties and its interactions with the series elasticity (Zajac, 1989; Todorov, 2003), but these properties are known to lose predictive value during locomotive situations where non-isometric and non-isotonic conditions that lead to transient and non-steady perturbation responses are commonplace (Lee et al., 2013; Sandercock and Heckman, 1997; Dick et al., 2017; Rice et al., 2020; Perreault et al., 2003, figure 1b). Furthermore, parameters in Hill-type models do not lend themselves to interpretation in terms of known mechanochemical processes within muscle. Despite these shortcomings, Hill-type models remain the most viable means to perform large-scale biomechanical simulations and optimal control calculations because of their low computational burden and ease of implementation (De Groote and Falisse, 2021; O’Neill et al., 2013; Miller, 2014).

The flip side to phenomenological models are those that incorporate the current state of understanding of the microscopic mechanochemical cycles. The hydrolysis of ATP by myosin II motors during formation of actin-myosin crossbridges underlie active muscle contraction and perturbation response (Lymn and Taylor, 1971; Holmes, 1997; Huxley and Simmons, 1971). Each myosin undergoes the biochemical Lymn-Taylor cycle in which it attaches to and detaches from actin filaments based on ATP capture, hydrolysis, and release of byproducts (Lymn and Taylor, 1971). There are different modeling approximations of this cycle that incorporate different numbers of intermediate states (figure 1b). The biochemical Lymn-Taylor cycle is coupled to a mechanical cycle that converts chemical energy to mechanical work. When attached, each myosin motor pulls on the actin filament to generate piconewton forces and around 10 nm power strokes. The collective action of many motors is modeled as an ensemble of stochastically cycling crossbridges whose force contributions add up (Huxley, 1957; Huxley and Simmons, 1971). Mechanical perturbation of the whole muscle ultimately perturbs the mechanochemical crossbridge cycle whose dynamics are load and strain-dependent (Palmer, 2010; Liu et al., 2018), which in turn alters the force produced by the ensemble and, in principle, manifests as bulk rheological properties of muscle. But crossbridge models employ numerous biochemical parameters that cannot be directly measured because of which we lack a mechanistic understanding of muscle’s emergent rheology. This is the so called scale-connection problem.

Addressing the scale-connection problem would overcome many of the shortcomings of both microscopic and macroscopic models, and is a major objective of neuromuscular research (Nishikawa et al., 2018). The distribution-moment formalism was a crucial step in on-going efforts to address it and presented a general approach to simplifying microscopic models without relying on the exact choice of how a crossbridge cycle is modeled (Zahalak, 1981, 1986; Zahalak and Ma, 1990). Specifically, the formalism assumed a Gaussian distribution for the fraction of bound crossbridges as a function of crossbridge strain such that the governing ensemble dynamics is approximated by the dynamics of measurable macroscopic quantities: stiffness, forces, and elastic energy storage. Thus, it is an approximate yet tractable mapping from these macroscopic quantities to the ensemble dynamics. But in assuming a preset distribution and not one that emerges from the crossbridge dynamics, it stops short of connecting scales back to the crossbridge cycle. Shortcomings of this approximation are evident from multiple perturbation experiments which show that the distribution-moment model does not accurately capture muscle forces, but more complex cross-bridge models may be able to do so by sacrificing computational speed (Cole et al., 1996; van den Bogert et al., 1998). So a gap remains in mapping crossbridge theory to the emergent rheology of its ensemble.

In this paper, we connect the dynamics of a single crossbridge to the rheology of an ensemble or population of crossbridges that represents a whole sarcomere. In particular, we analyze the ensemble behavior of several different crossbridge models to identify a minimal parameter-set that affects their emergent rheology. The paper begins with a brief tutorial on rheological characterization of materials using complex modulus and dynamic stiffness. Then we use numerical simulations of large ensembles of crossbridges, ranging in complexity from two-state to five-state crossbridge models, to identify commonalities and differences in their perturbation responses. Next we analytically derive the rheology of a two-state crossbridge model and show how a single exponential relaxation process with one time-constant suffices to capture its rheology, how that relaxation is related to the crossbridge dynamics, and provide a mechanical interpretation in terms of elastic and viscous material properties. We then extend the characterization to more complex crossbridge models by adding multiple exponential relaxations and discuss the implications of our work towards the development of computationally efficient and parametrically parsimonious models that are still interpretable in terms of the microscopic mechanochemical dynamics of actomyosin crossbridges.

## Preliminaries

We focus here on the linear rheological characterization of muscle, which has proven useful in developing predictive models of muscle behavior (Rack and Westbury, 1974; Nguyen et al., 2018; Bizzi et al., 1982) and in interpreting *in vivo* muscle function for both large and small perturbations (Palmer et al., 2020; Niederer et al., 2019). A central quantity in linear rheology is the material modulus defined as the ratio of the measured stress to the applied strain. This modulus for muscle depends on the rate at which the strain is applied and varies over time. Using the Laplace or Fourier transform (Ogata, 2004), the applied strain and the resulting stress response are each decomposed into a sum of sinusoids of different temporal frequencies, and their ratio yields a frequency-dependent modulus. This characterization, a core tool in oscillatory rheology (Weitz et al., 2007; Tschoegl, 2012), provides an interpretation of muscle material properties in terms of frequency-dependent loss, storage, and complex moduli (Kawai and Brandt, 1980; Nguyen et al., 2018; Nguyen and Venkadesan, 2021).

If the applied strain acting as an external perturbation is a small-amplitude sinusoidal wave, then the resulting stress perturbation respones is approximately sinusoidal with both in-phase and out-of-phase components with respect to the applied strain (figure 2(a-b)). The amplitude of the in-phase stress and out-of-phase stress divided by the applied strain amplitude define the storage modulus *E*^*′*^(*ω*) and loss modulus *E*^*′′*^(*ω*), respectively, and are functions of the oscillatory frequency *ω*. The complex modulus *E*^*∗*^(*ω*) = *E*^*′*^(*ω*) + *iE*^*′′*^(*ω*) is a compact representation of the spring-like storage modulus and damper-like loss modulus. The dynamic stiffness *K*(*ω*) of sinusoidal analysis is related to the complex modulus as *K*(*ω*) = (*A*/*L*_0_)|*E*^*∗*^(*ω*)|for a muscle tissue of cross-sectional area *A* and length *L*_0_. If muscle were elastic like a Hookean spring, then *K*(*ω*) is a constant independent of frequency. If muscle were viscous like a Newtonian fluid, then *K*(*ω*) is linearly proportional to frequency. In reality, muscle is neither extreme and exhibits intermediate viscoelastic behavior (figure 2c).

**Figure 2:**
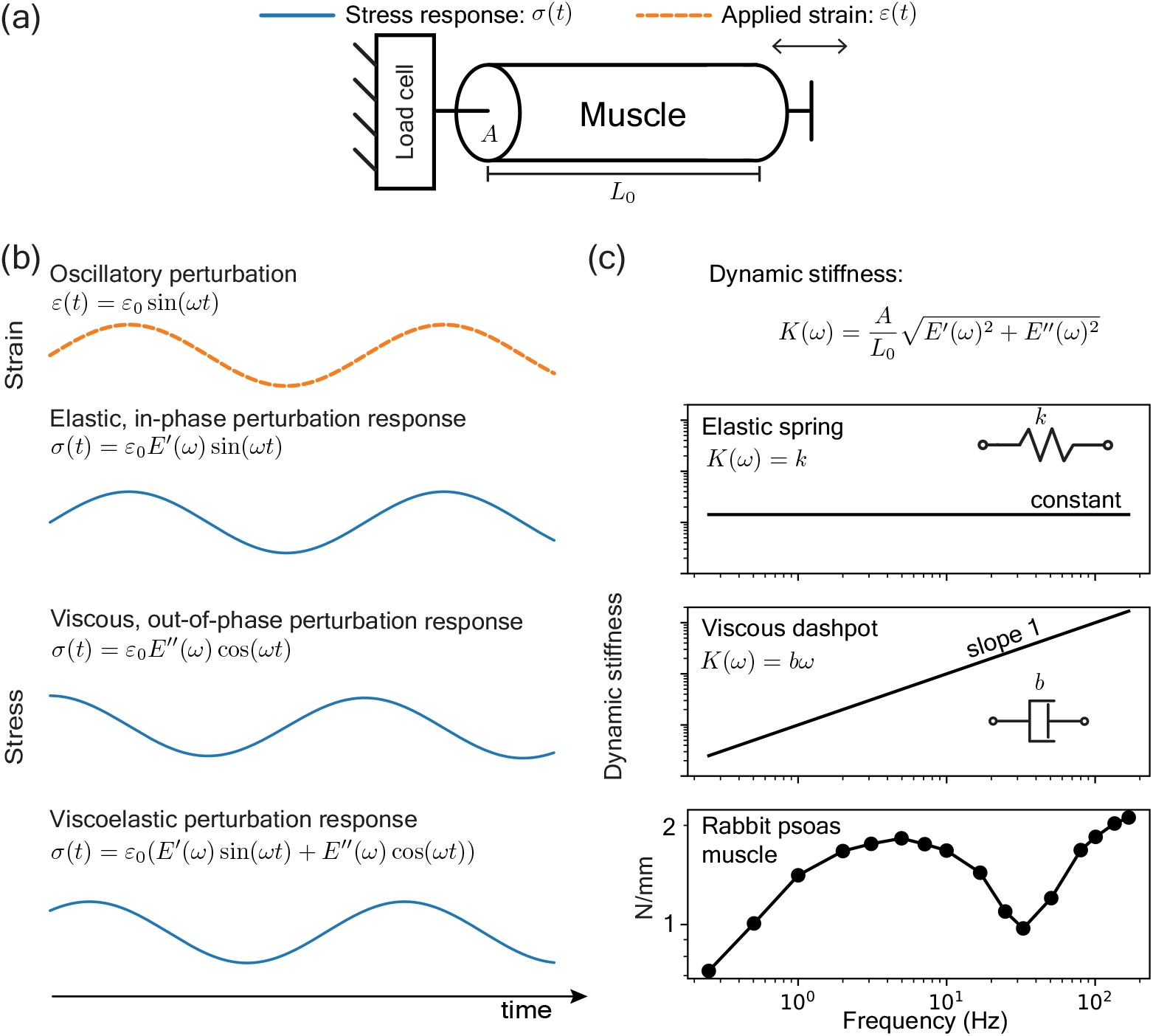
Linear rheological characterization of muscles. (a) Schematic of an experimental setup to measure a muscle’s response to perturbations. (b) For an oscillatory length perturbation, the linear rheological response is characterized by an elastic modulus *E*^*′*^(*ω*) that measures the in-phase response and a loss modulus *E*^*′′*^(*ω*) that measures the out-of-phase response. A general viscoelastic response has both in-phase and out-of-phase components. (c) The dynamic stiffness *K*(*ω*) is a third oscillatory rheological characterization. It is constant if the muscle is elastic and linearly proportional to frequency if the muscle is viscous. The dynamic stiffness of a rabbit psoas muscle is viscoelastic with features more complicated than elastic springs and viscous dashpots alone. The rabbit psoas plot is adapted from figure 3 of Kawai and Brandt 1980.

## Results

### Numerical simulations of ensemble crossbridge models

We numerically compute the emergent rheology of an ensemble or population of independently cycling crossbridges using previously published crossbridge models. The aim here is to survey the literature and use models that differ in complexity and in the types of muscle phenomena that each were designed to capture. To that end, we implement four crossbridge models ranging from two to five internal states (Huxley, 1957; Murase et al., 1986; Smith, 1998; Lombardi and Piazzesi, 1990). Importantly, our selection of crossbridge models is a representative but not exhaustive survey of all muscle phenomena under active research. For example, the super relaxed myosin state introduces a new detached state with orders of magnitude lower ATP turnover than the normal detached state (Stewart et al., 2010; McNamara et al., 2015). Thin filament activation and cooperative mechanisms can introduce a coupling between crossbridges that break the assumption of independently cycling crossbridges (Walcott, 2014). Spatially explicit sarcomere models account for the distribution of crossbridges within the sarcomere and thus also for position-dependent crossbridge dynamics (Daniel et al., 1998; Williams et al., 2012; Kosta et al., 2021). Although the crossbridge literature is broader than our selection, or any similarly sized selection can reasonably sample, our objective with numerical simulations is to connect the emergent ensemble rheology with the detailed modeling choices regardless of the specific crossbridge model.

The simplest of the crossbridge models considered is Huxley’s original two-state formulation that helped set the foundation for the whole class of crossbridge models developed since and its emergent rheology can thus serve as a point of reference (Huxley, 1957). In it, Huxley idealizes crossbridges as molecular springs that stochastically attach to and detach from a rigid thick and thin filament backbone. The total elastic resistance of a population of these crossbridges is then taken to characterize the active and transient forces of muscles and, in turn, also the rheology of muscles. The other more complex crossbridge models refine Huxley’s formation by incorporating additional attached and detached states which allows for new and higher-order dynamics that Huxley’s two-state model may not capture. The three-state model by Murase et al., 1986 (Murase et al., 1986) showed that a second attached internal state is necessary to capture the three dominant sinusoidal processes observed in insect flight muscles. The four-state model by Smith, 1998 (Smith, 1998) proposed a minimal kinetic scheme to capture the force transients resulting from the phosphate release and ATP capture of the Lymn-Taylor cycle (Lindstedt et al., 2001). And lastly, the five-state model of Lombardi and Piazzesi, 1990 (Lombardi and Piazzesi, 1990) proposed a relatively complex kinetic scheme to capture the force transients of a frog skeletal muscle fiber under steady lengthening that, notedly, incorporated a forced detached state accessible only when crossbridges are stretched beyond a critical distance. The numbers of parameters involved for each model are many with 4 for Huxley’s two-state model, 16 for the three-state model, 17 for the four-state model, and 24 for the five-state model. These parameters define the neutral length, stiffness of each attached crossbridge state, and the transition rates between them. A natural question that arises is how the emergent ensemble rheology depends on these parameters and, given such a vast parameter space, whether there is parameter-reduction that recapitulates the emergent rheology. We shall explore if such a parameter reduction is possible in the numerical simulations below.

The emergent ensemble rheologies of the selected crossbridge models are computed in terms of the dynamic stiffness, the variation of which with frequency determines whether the ensemble is like an elastic solid or a viscous damper. Numerically, we integrate each model’s governing differential equations to compute the force response of an ensemble of crossbridges to a step length perturbation. The ensemble is initially at steady-state equilibrium so the emergent rheology arises from the ensemble’s relaxation back to steady-state. Transforming the computed force response and the imposed step length perturbation into the frequency domain and taking their ratio directly yields the dynamic stiffness (Methods A).

We find that a simple behavior emerges at high and low frequencies regardless of model complexity and that the difference between crossbridge models manifest only on intermediate frequencies (figure 3b). Specifically, the dynamic stiffness is constant at high frequencies and linearly proportional to *ω* at low frequencies, i.e. the dynamic stiffness is approximately

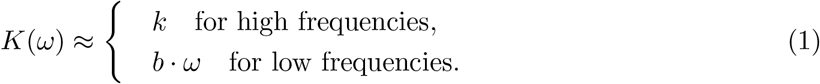

**Figure 3:**
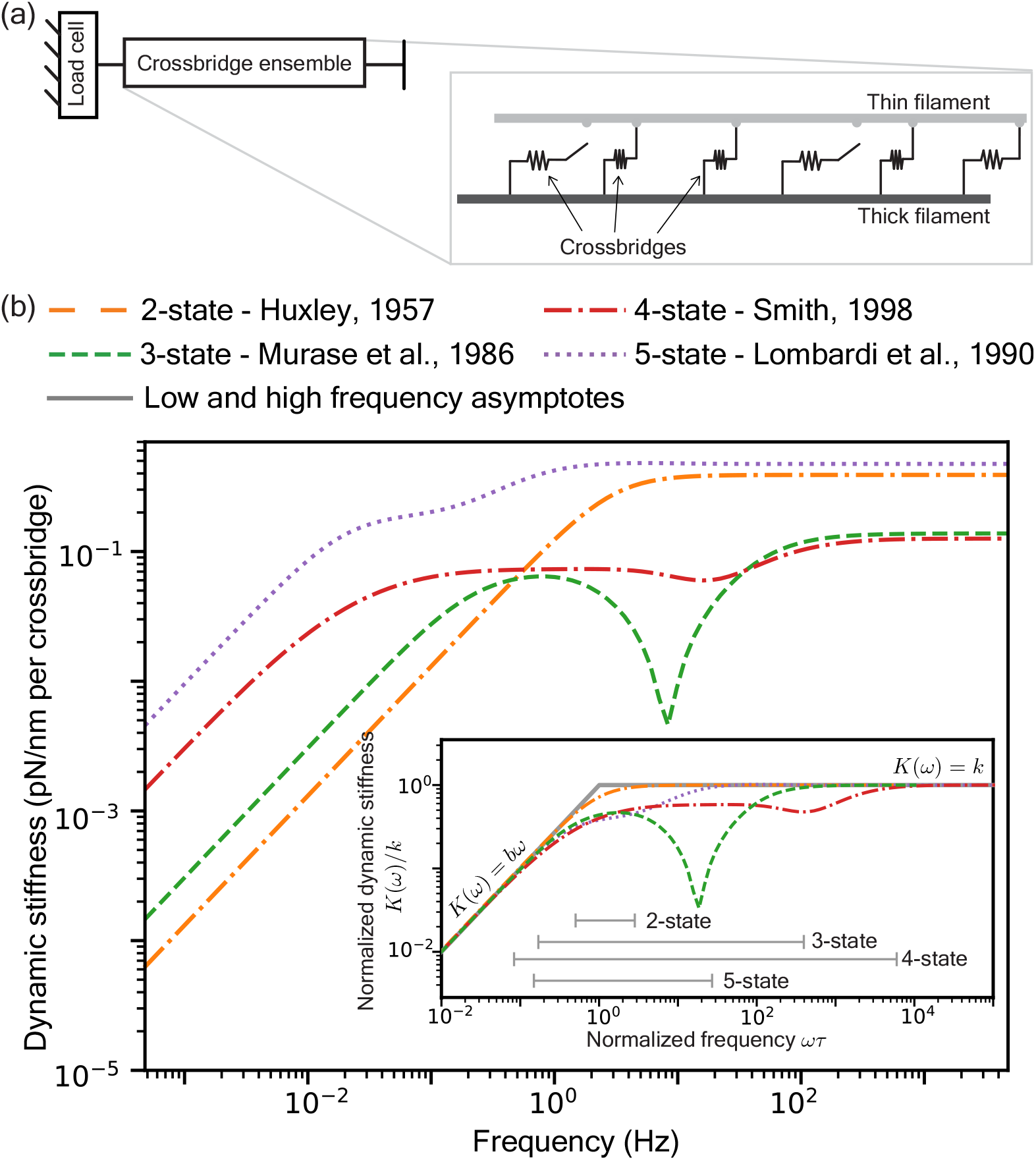
Rheology of the ensemble dynamics of different crossbridge models. (a) An ensemble of crossbridges operate collectively to model muscle forces. (b) The frequency-dependent dynamic stiffness of different crossbridge models resembles an elastic spring at high frequencies and a viscous dashpot at low frequencies. (inset) Collapse of different models onto equation (1) by normalizing the axes. For each model, the dynamic stiffness is normalized by the value *k* computed at the highest frequency. The frequency axis is normalized by a time-constant *τ* at which the two asymptotic behaviors match. The gray bars indicate the bandwidth over which the models deviate by more than 5% from their asymptotic behavior s.

These asymptotic regimes imply that the ensemble is an elastic solid with spring constant *K* at high frequencies and a viscous damper with damping coefficient *b* at low frequencies. But the dynamic stiffness at intermediate frequencies is more complicated in their frequency dependence, differs between the model variants, and does not immediately present a mechanical interpretation. We calculate a bandwidth in this intermediate frequency range where the dynamic stiffness differs by more than 5% from the asymptotic response (grey bars in inset of figure 3b) as a measure of complexity in the emergent rheology. The bandwidth is also indicative of computational cost because the largest time-step that can be used for numerical integration is governed by the highfrequency end of the bandwidth whereas the duration of time needed for the transient response to settle down is governed by the low-frequency end. We find that the two-state Huxley model has the smallest bandwidth and is followed by the five-state, three-state, and four-state models. This ordering does not follow the number of internal states so the complexity of the emergent rheology does not correlate with complexity of the crossbridge model. For example, although the five-state model requires 24 parameters, the largest amongst all models considered, its emergent rheology is well-approximated as a high-frequency elastic spring and a low-frequency vicious damper which requires only two parameters and suggests that a drastic parameter reduction from the detailed crossbridge model to its emergent rheology is possible. We shall analyze the mathematical forms of these crossbridge models to derive the exact form of the parameter reduction.

### Rheology of the generalized two-state crossbridge model

We analytically derive the rheology for the two-state crossbridge ensemble to examine the rheological features at intermediate frequencies that are not captured by the high and low frequency asymptotes (equation (1)). We refer to (Smith, 1998) for a conceptually identical but more general derivation of a four-state crossbridge model while we focus here on the simpler two-state variant to demonstrate how modeling choices lead to different rheological features.

Two-state crossbridges cycle between making and breaking bonds between rigid thick and thin filaments. The attachment rate *f*(*x*) and detachment rate *g*(*x*), to form and break bonds, respectively, are functions of the displacement *x* between the location of the myosin motor on the thick filament and the nearest binding site on the thin filament. At every displacement *x* and time *t*, the proportion of attached crossbridges is *n*(*x, t*), also known as the bond distribution, and the detached proportion is (1 −*n*(*x, t*)). The bond distribution *n*(*x, t*) evolves in time according to

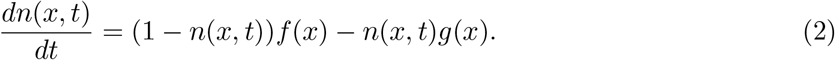

This first-order differential equation has a steady-state bond distribution *n*_ss_(*x*) = *f*(*x*)/(*f*(*x*) + *g*(*x*)) at equilibrium. Starting with this equilibrium bond distribution, a step length perturbation of amplitude *a* is delivered at *t* = 0 to elongate the whole system and hold it there. Every crossbridge that was a displacement *x* away from its nearest binding site is now a displacement (*x*+*a*) away and the distribution of crossbridges *n*(*x, t*) evolves in time to relax back to its equilibrium state according to equation (2). The response to this step length perturbation is obtained using *n*(*x*, 0) = *n*_ss_(*x*−*a*) as the initial condition and integrating equation (2) to find

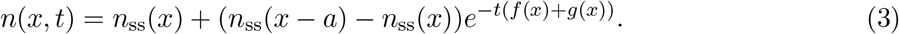

Therefore, *n*(*x, t*) exponentially relaxes to its steady-state *n*_ss_(*x*) at a rate *f*(*x*)+*g*(*x*). We illustrate this relaxation process in figure 4a using Huxley’s rate functions (Huxley, 1957). The force Δ*F*(*t*) in response to the step length perturbation is determined by the difference between *n*(*x, t*) and its steady state *n*_ss_(*x*). In terms of crossbridge stiffness λ_xb_ in units of [force/length] and crossbridge density *M* in units of [length]^*−*1^, the force response is

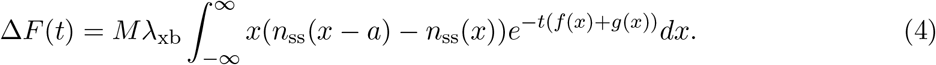

**Figure 4:**
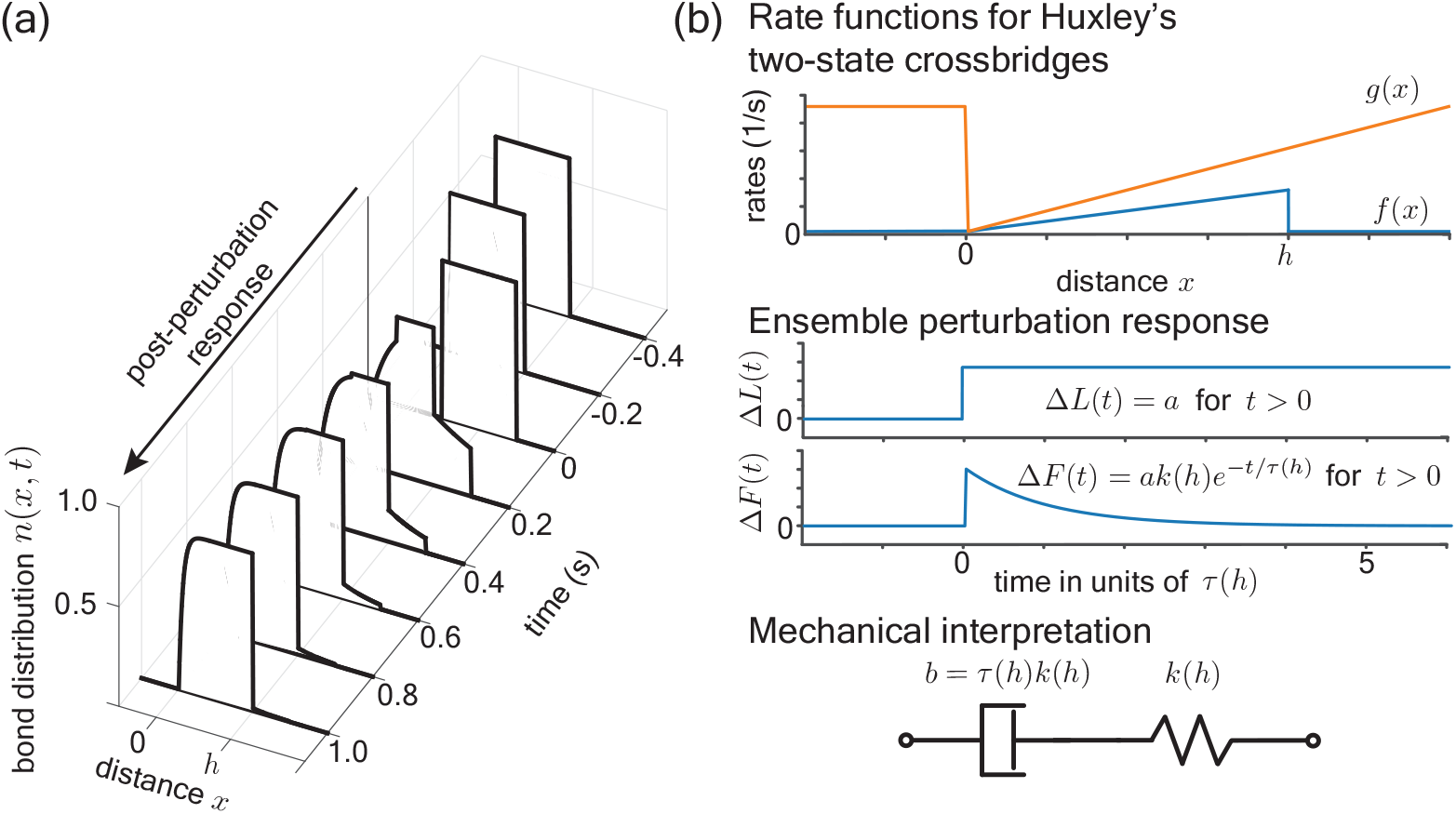
The rheology of an ensemble of Huxley’s two-state crossbridges. (a) The time-evolution of the bond distribution in response to a step length perturbation. (b) Rheology of an ensemble of Huxley’s two-state crossbridges. The ensemble’s perturbation response exponentially relax with time-constant *τ*(*h*) = 1/(*f*(*h*) + *g*(*h*)) where *f*(*x*) and *g*(*x*) are the rate functions and *h* is the powerstroke distance. A mechanical interpretation of this single exponential relaxation is of a spring in series with a dashpot.

By restricting our attention to the linear regime, *i*.*e*. a small perturbation *a*, the force response simplifies to

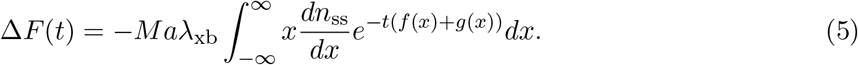

This perturbation response may be rewritten as a sum of exponential relaxations with a displacement-dependent relaxation time-constant *τ*(*x*) and displacement-dependent ensemble stiffness *k*(*x*) that is given by,

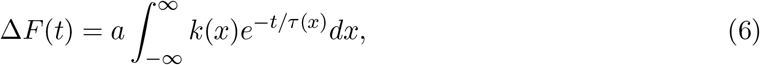

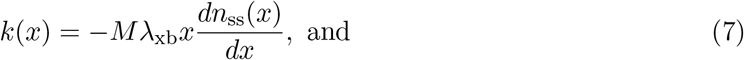

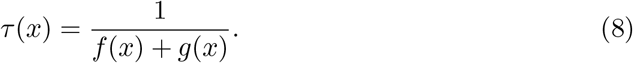

Therefore, the entire relaxation process may be thought of as being comprised of an infinity of exponentially relaxing sub-processes where the displacement-dependent stiffness *k*(*x*) is the weight and *τ*(*x*) the relaxation timescale for each sub-processes. To obtain a dynamic stiffness *K*(*ω*), we use the Laplace transforms of equation (6) and of the step length perturbation to arrive at

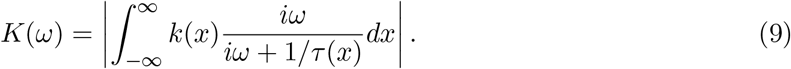

The integrand is the mechanical impedance of a Maxwell body, a spring mechanically in series with a dashpot, and a mechanical analogy can thus be drawn between ensemble crossbridge dynamics and Maxwell bodies. We demonstrate the analogy for the two-state model using Huxley’s choices for the rate functions *f*(*x*) and *g*(*x*) (Huxley, 1957, figure 4b) given as

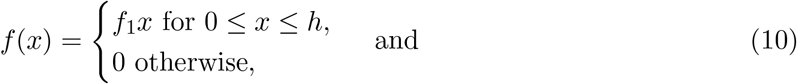

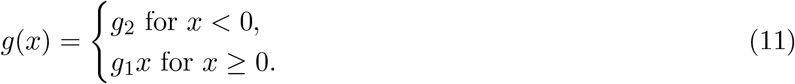

which leads to the steady-state bond distribution *n*_ss_(*x*)

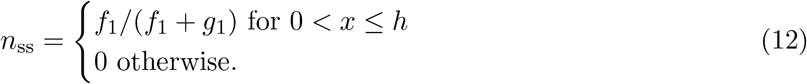

Applying equations (7 – 9), the dynamic stiffness corresponding to Huxley’s choices of *f*(*x*) and *g*(*x*) is that of a single Maxwell body with stiffness *k*(*h*) and damping coefficient *b* = *k*(*h*)/(*f*(*h*)+*g*(*h*)) = *k*(*h*)*τ*(*h*). To add more detail, the rectangular waveform of the steady-state bond distribution (equation (12)) implies that *k*(*x*) of equation (7) is a sum of two *δ*-functions, a negative one at *x* = 0 and a positive one at *x* = *h*. However, because the delta at *x* = 0 does not contribute to the perturbation response, only the delta at *x* = *h* remains and the overall perturbation response is that of a single Maxwell body. Huxley’s choices thus lead to an exact correspondence between ensemble crossbridge dynamics and a single Maxwell body, which justifies the mechanical interpretation for the high and low frequency rheological behaviors observed in figure 3 as elastic and viscous behaviors. The spring-like behavior at high frequencies is not surprising because springs are built into ensemble crossbridge models by idealizing each crossbridge as an individual spring, all of which adds up at high frequencies. On the other hand, there are no dampers built into crossbridge models and the low-frequency behavior thus arise as an emergent property of the ensemble rather than as an intrinsic property of crossbridges. Specifically, each crossbridge dissipate stored elastic energy as they cycle and this dissipation averaged over the entire ensemble and over many cycles manifests as an effective damping coefficient at low frequencies.

The stiffness distribution *k*(*x*) of equation (7) is well-approximated by a *δ*-function for Huxley’s two-state model leading to single exponentially relaxing sub-process that dominates the entire perturbation response. Thus whether we need the full ensemble model or if we can approximate the response with just a few discrete relaxation sub-processes depends on whether *k*(*x*) shows localizations in *x* or not. We hypothesize that, for all crossbridge models considered in this paper, the localization is a dominant feature and that only a few discrete relaxation sub-processes are needed to capture the emergent rheological features on intermediate frequencies (figure 3b). Because *k*(*x*) is proportional to the derivative of the steady-state bond distribution, its localization and the resulting exponential sub-process directly equate to either a sharp rising or falling edge of the steady-state bond distribution, or of multiple bond distributions for crossbridge models with multiple attached states. We shall verify the hypothesis by fitting exponential sub-processes to the dynamic stiffness computed for the crossbridge models and examine if those fitted processes corresponds to rising or falling edges of the bond distribution.

### Low-order models of multi-state crossbridge ensembles

Crossbridge dynamics are generally more complex than Huxley’s two-state model and may have multiple internal states that introduce higher-order dynamics. Nevertheless, the perturbation force response initially arises from stretching bound crossbridges and its long-term relaxation is because of a redistribution of the crossbridges between the various possible states. This suggests an approach to tackle the behavior in the intermediate frequencies where crossbridge models were shown to deviate from the simple asymptotic behavior of a Maxwell body (figure 3b). Namely that, under more complicated attachment-detachment dynamics, the perturbation response would show multiple exponential relaxation sub-processes. So we investigate whether a finite number of exponential relaxations can capture the emergent ensemble rheologies in spite of possible complexities, under the hypothesis that a few sub-processes dominate the response. In other words, we examine whether the ensemble’s frequency-dependent dynamic stiffness *K*(*ω*) can be approximated by a sum of *N* Maxwell bodies according to,

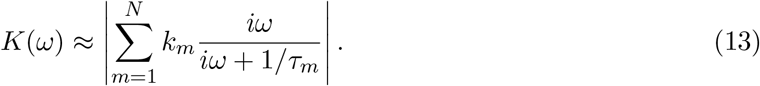

We find that at most three exponentially relaxing sub-processes, i.e. *N* = 3 in equation (13), are needed for any of the crossbridge models to accurately capture the frequency-dependent stiffness over seven orders of magnitude on the frequency axis (figure 5a, Methods B, Figure 5 - Table Supplement 1). Huxley’s two state model fit exactly with a single sub-process, consistent with our earlier derivation. The five-state model is also well-approximated by two sub-processes. The three- and four-state models are mostly fitted by three sub-processes. By encapsulating the emergent rheology using few sub-processes, the number of parameters associated with the crossbridge models are fewer by 2–6 fold, with the biggest gains in reducing parametric complexity for the more complex models with multiple internal states (table 1). Overall, the accurate fits demonstrates that, although crossbridge models incorporate a vast parameter space, the rheologies exhibited by their ensemble dynamics are far simpler and dominated by only a few exponentially relaxing sub-processes.

**Figure 5:**
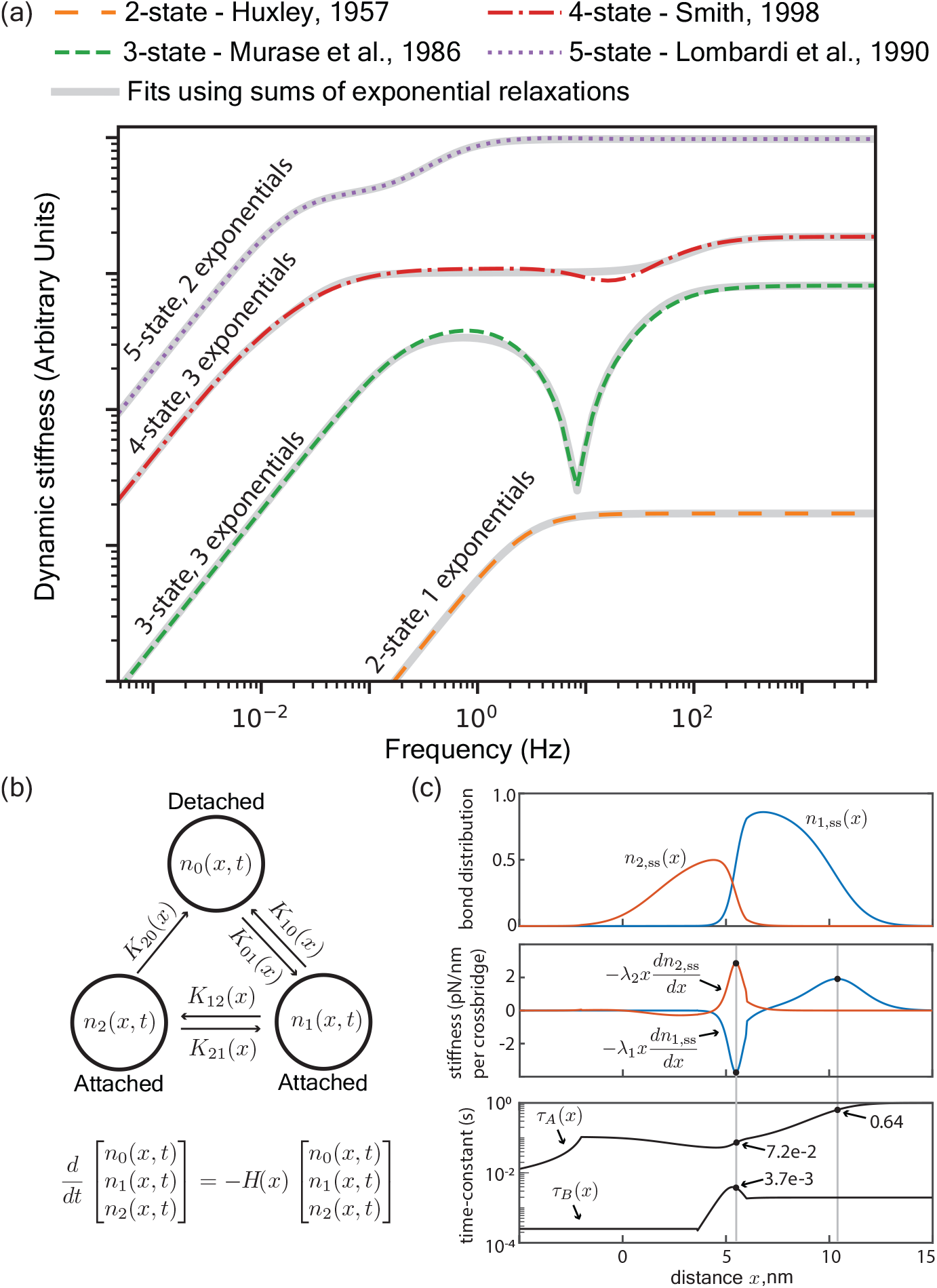
(a) The dynamic stiffness of different crosssbridge ensembles are accurately fitted by at most 3 time-constants, each associated with an exponential process (see Figure 5 - Table Supplement 1 for fitted values). The traces are vertically shifted for clarity (see Figure 3 - Figure Supplement 1 for an unmodified plot). (b) Kinetic scheme of a three-state crossbridge model (Murase et al., 1986). The matrix *H*(*x*) is a state transition matrix with two non-zero eigenvalues 1/*τ*_*A*_ and 1/*τ*_*B*_. (c) Examination of the derivatives of the steady-state distributions shows that the local peaks accurately identifies the time-constants fitted in Figure 5 - Table Supplement 1.

**Table 1:** Reduction in parametric complexity. Number of parameters in the full model versus the low-order approximation in terms of exponential sub-processes.

| Model | Full model | Low-order approximation |
| --- | --- | --- |
| Huxley, 1957 | 4 | 2 |
| Murase et al., 1986 | 16 | 6 |
| Smith, 1997 | 17 | 6 |
| Lombardi et al., 1990 | 24 | 4 |

The fitting process finds sub-processes acting on vastly different timescales and has also been previously applied to the describe the rheology of muscles in terms of sub-processes (Kawai and Brandt, 1980; Kawai et al., 1993). We next address the question of how to relate the fitted values to crossbridge dynamics. The analysis of Huxley’s two-state model provides a possible method to connect the scales from crossbridge dynamics to ensemble rheology. The dynamic stiffness of the entire ensemble depends on the integral over a continuum of exponential sub-processes *k*(*x*) *e*^*−t/τ*(*x*)^, with stiffness *k*(*x*) and time-constant *τ*(*x*) that depend on the crossbridge displacement *x*. Importantly, we found that sub-processes where the stiffness *k*(*x*) peaked dominated the ensemble rheology, and the associated relaxation time *τ*(*x*) governed the frequency-dependence of the ensemble stiffness *K*(*ω*).

We investigated whether that insight carries over to the three-state model (Murase et al., 1986). The three-state model serves as an intriguing test-case for several reasons. It required the most number of fitted exponential sub-processes. The three-state model also exhibits features that are most unlike the two-state model, including a significant dip in the dynamic stiffness that is indicative of a negative exponential sub-process that has been previously attributed to a delayed tension rise observed in a muscle’s perturbation response (Kawai et al., 2021). The three-state model has two attached states with steady-state distributions *n*_1,ss_(*x*) and *n*_2,ss_(*x*), according to the kinetics illustrated in figure 5b. We present here a simplified version of the analysis by assuming independence of the kinetic equations for each state, which allows us to identify separate sub-processes associated with each bound state. Methods C presents a full analysis without this simplification but shows nearly identical findings. To identify the dominant sub-processes of the three-state model, we examine the sub-process stiffness distributions associated with each of the two bound states, namely, *λ*_1_*xdn*_1,ss_(*x*)/*dx* and *λ*_2_*xdn*_2,ss_(*x*)/*dx* (figure 5c). There are four local extrema, two per attached state, of the sub-process stiffness distributions (figure 5c) of which three are clearly dominant. The fourth one appears near *x* = 3nm and is considerably smaller than the other three by an order of magnitude. So we estimate the time constant *τ* associated with the three dominant local extrema of these sub-process stiffness distributions and find that they resemble the three fitted time-constants (Figure 5 - Table Supplement 2).

This level of correspondence is surprising because, generally, the sum of exponentials is not an exponential. Being able to approximate the ensemble behavior as a sum of exponentials implies that we are able to represent an integral over exponentials as a discrete sum of few exponentials. That the sub-process stiffness distribution has just a few extrema is part of the reason, but additionally, the time-constants should not vary too rapidly in neighborhood of the extrema for the approximation to hold. Therefore, the surprising, but simplifying finding of the analysis is that the governing equations of two, three, four, and five state crossbridge models are such that the integral over a continuum of exponential sub-processes is well approximated by a sum over few discrete exponential sub-processes.

Our examination of multi-state crossbridge models puts forth the following connection between the exponential relaxation fitted to muscle rheology and the dynamics of crossbridge cycling. Namely, each fitted relaxation results from either a sharp rising or falling edge of the steady-state bond distribution of attached crossbridge states. Assuming that most crossbridges attach at positive *x*, the fitted stiffness is positive for a falling edge and negative for a rising edge. The fitted time-constant are associated with the attachment and detachment rates corresponding to the rising or falling edges of the steady-state bond distributions. Furthermore, because the forces arising from the edges are modulated by *x*, the falling edge at larger *x* can dominate over the rising edge at smaller *x*, such as in the case of Huxley’s two-state model. Therefore, although the bond distributions themselves may be of a complicated functional form due to the vast parameter space that multi-state crossbridge models tend to require, the emergent ensemble rheology is simpler and dominated by few rising and falling edges. This simplification affords a parameter reduction of multi-state crossbridge models down to two per exponential relaxation, a stiffness and a time-constant, and will allow us to compute the rheological behavior of ensemble crossbridge models in large-scale musculoskeletal simulations in a manner as efficient as phenomenological Hill-type muscle models but without compromising on the mechanistic, crossbridge-based understanding.

## Discussion

Our results simplify the complexity of crossbridge models and show how lower-order dynamics emerge from the connection across vastly different scales of crossbridge dynamics and ensemble rheology. These results are similar to Zahalak’s distribution-moment formalism (Zahalak, 1981), but go further and analyze how modeling choices of a single crossbridge cycle give rise to macroscopically measurable rheological properties. We focused on the dynamic stiffness which generalizes notions of elastic stiffness and viscous damping into a viscoelastic property that depend on the time-duration of interest. Specifically, we showed that the emergent rheology of crossbridge ensembles can be accurately fitted by only a few exponential sub-processes. Such fitting procedures have been previously applied to empirical muscle data to identify the dominant exponential sub-processes (Kawai and Brandt, 1980; Kawai et al., 1993), and our findings add the understanding that these sub-processes correspond to rising or falling edges of the steady-state bond distributions.

Phenomenological models, with few and experimentally measurable parameters, and lesser computational overhead, present numerous advantages over detailed multiscale and spatially explicit models (Kosta et al., 2021). Experimental measurability and the computational efficiency of Hill-type models over PDE descriptions of crossbridge ensembles is a principal reason for the widespread adoption of Hill-type muscle models (De Groote and Falisse, 2021; O’Neill et al., 2013; Miller, 2014) despite their shortcomings (Nishikawa et al., 2018; Nishikawa and Huck, 2021). Therefore, augmenting Hill-type models with experimentally measurable exponential sub-processes will facilitate large-scale biomechanical simulations without sacrificing the grounding of these muscle models in crossbridge theory. But the finding that the rheological response of mean field models of cross-bridge ensembles simplifies to sums of exponentials identifies some fundamental gaps in current understanding of muscle’s response to perturbations. We discuss here two main issues brought to light by the analyses.

One limitation is that systems with exponential relaxations have limited memory determined by its time-constant. So an ensemble of crossbridges cannot remember its perturbation history far beyond its longest time-constant. Residual force enhancement and force depression are history-dependent phenomena that have implications for the motor control of muscles (Herzog, 2004), and are in clear contradiction with having an exponentially decaying memory. In these phenomena, the forces exhibited by a muscle fiber upon being stretched or shortened to a final length relax towards a value that is persistently higher (force enhancement) or lower (force depression) than if the muscle fiber were isometrically held at the final length. So, unlike exponential relaxation, muscle exhibits long-lived memory of being stretched or shortened. This suggests that history-dependent muscle phenomena are outside the purview of current crossbridge models. However, such history-dependence parallels phenomena found in other biological and non-biological materials for which a phenomenological fractional viscoelastic model captures power-law or non-exponential relaxation (Jaishankar and McKinley, 2013; Bagley and Torvik, 1983; Kollmannsberger and Fabry, 2011). Deriving inspiration from these fractional viscoelastic materials may help to better understand the physical underpinnings of residual force enhancement and depression in muscles. Nevertheless, our analyses of multiple crossbridge models show that they are not capable of exhibiting fractional viscoelastic or other non-exponential responses over long time durations.

A second limitation is the neural control of muscle’s relaxation timescales (Nguyen et al., 2018). It is common experience that muscle can remain stiff and behave like an elastic solid over long timescales when it is highly stimulated by the nervous system; consider holding the elbow at a fixed posture. On the flip side, when muscle is held in a relaxed state, stresses can dissipate over much shorter period; consider throwing a fast ball where the biceps stretch rapidly. Contrary to such common experience, we find that the time-constants arising from crossbridge ensembles do not depend on the number of engaged crossbridges, i.e. not dependent on neural drive. The number of binding sites available for crossbridges increases with calcium concentration. However, more crossbridges would only increase the sub-process stiffness *k* and leave the relaxation time-constant *τ* unchanged. This is because, in all the models we investigated, crossbridge cycling dynamics are independent of the total number of crossbridges. Some crossbridge models incorporate faster crossbridge cycling with higher calcium (Zahalak, 1986; Zahalak and Ma, 1990; Walcott, 2014). While this leads to higher contractile forces, the time-constant would in fact decrease with higher calcium rather than increase. Therefore, crossbridge models cannot simultaneously capture higher contractile force and slower stress relaxation when the neural drive to muscle is greater.

The mismatches between exponential relaxations, known muscle phenomena, and requirements of neural control all suggest that either crossbridge models currently lack an essential feature or that non-crossbridge elements come into play. There are multiple possible candidate resolutions that we speculate may operate in unison, and certainly require further experimental investigation to improve our understanding of muscle. Thick and thin filament compliance between neighboring crossbridges can store elastic energy that dissipate far slower than the crossbridges themselves and affect muscle forces on long experimental timescales (Mijailovich et al., 1996; Daniel et al., 1998). Intersarcomeric dynamics may lead to dynamics not captured by single sarcomere model such that the number of participating sarcomeres can drastically alter the rheology of a muscle fiber (Shimamoto et al., 2009; Campbell, 2009; Caruel and Truskinovsky, 2018). Titin could be another factor that can directly modulate force transmission across a sarcomere and therefore affect crossbridge and inter-sarcomere dynamics (Nishikawa et al., 2012; Powers et al., 2014). Neural feedback and reflexes could introduce additional time-constants depending on motor tasks, but is limited by transmission delays on the order of 50ms for humans (Johansson and Cole, 1994; Hogan et al., 1987). These and other possibilities require further investigation, but point to a shortcoming in the current crossbridge theory for muscle forces and rheology.

## Author Contributions

K.D.N. and M.V. conceived the research, designed the research, analyzed and interpreted the results, and wrote the paper.

## Methods

### A Dynamic stiffness calculation of crossbridge models

We detail here the numerical calculation of dynamic stiffness for ensemble crossbridge models. Although each model differs in number of internal states, a general differential equation for the mass balance between states can be written. Let *j* be the index for the states, then *n*_*j*_(*x, t*) is the distribution of crossbridges in the *j*^*th*^ state at time *t* and with displacement *x* to the nearest binding site. The mass balance will take the general form

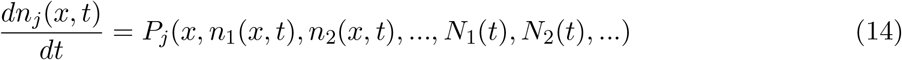

where 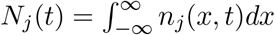 is the total proportion of crossbridges in state *j* and *P*_*j*_ is a function that depends on the crossbridge model. We use *P*_*j*_ as given by four different models varying from two to five internal states (Huxley, 1957; Smith, 1998; Murase et al., 1986; Lombardi and Piazzesi, 1990).

We impose a step length perturbation of amplitude *a* and use the output force perturbation response to numerically compute each model’s dynamic stiffness. Specifically, we numerically integrate equation (14) subjected to the initial condition *n*_*j*_(*x*,0) = *n*_*j*,ss_(*x* −*a*) using the explicit fourth-order Runge-Kutta (RK4) scheme (Press et al., 2007). The integration takes time steps of size *δt* and terminates at a finite time *T*. The output force perturbation response is a discrete time series Δ*F*_*m*_ indexed in time by *m* and calculated as

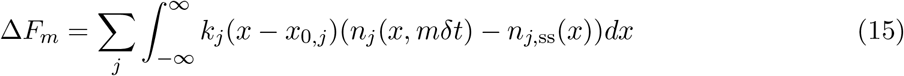

for stiffness *K*_*j*_ and neutral length *x*_0,*j*_ that depends on the *j*^*th*^ crossbridge state. The dynamic stiffness *K*(*ω*) is then computed in terms of z-transforms as

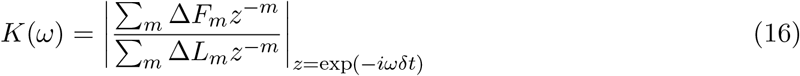

where Δ*L*_*m*_ is the step length perturbation equal to *a* for all *m* ≥0 and zero otherwise.

To calculate the dynamic stiffness for Huxley’s two-state model (Huxley, 1957) in dimensional units, we use parameter values (*f*_1_,*g*_1_,*g*_2_) = (15, 4, 85)s^−1^provided by (Zahalak and Ma, 1990) (Zahalak and Ma, 1990). We also used a crossbridge stiffness *K* _*x* b_= 0.5pN/nm that is consistent with literature values (Lombardi and Piazzesi, 1990; Smith, 1998).

### B Fitting procedure

We detail here the process of fitting exponential relaxations to the dynamic stiffness of different crossbridge models (figure 3d). We denote *ω*_*j*_ as the frequency indexed by *j* and logarithmically sampled from 5 ∗ 10^−4^Hz to 5 ∗ 10^3^Hz. We also denote *K*(*ω*_*j*_) as the numerically determined dynamic stiffness at frequency *ω*_*j*_. The objective is to find N-pairs of parameters (*K*_*m*_, *τ*_*m*_) indexed by m such that the function

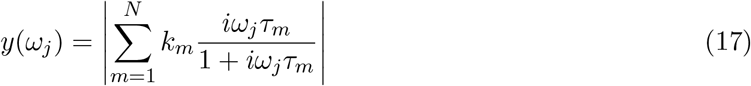

accurately captures *K*(*ω*_*j*_) where *N* is the number of exponential relaxations to fit and *i* is the imaginary number. We defined fit to be the N-pairs of parameters (*k*_*j*_, *τ*_*j*_) that minimizes the cost

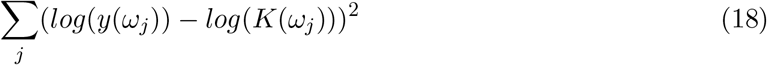

where we use the log function to equally weight behaviors at vastly different frequencies. The fit was performed starting at *N* = 1 and *N* is incrementally increased until there is no appreciable difference between the fits at *N* and at *N* + 1.

All optimization procedures are performed using the Python ‘scipy.optimize’ library (Virtanen et al., 2020).

**Table Figure 5 - Table Supplement 1:** Fitted parameters for figure 5a. Stiffness values are in units of pN/nm per crossbridge and time-constants are in units of seconds.

| Model | $k_1$ | $\tau_1$ | $k_2$ | $\tau_2$ | $k_3$ | $\tau_3$ |
| --- | --- | --- | --- | --- | --- | --- |
| Huxley, 1957 | 0.39 | 5.3e-2 | - | - | - | - |
| Murase et al., 1986 | 0.09 | 0.59 | -0.10 | 6.8e-2 | 0.15 | 3.2e-3 |
| Smith, 1997 | 0.05 | 2.0 | -0.12 | 3.6 | 0.20 | 1.4e-3 |
| Lombardi et al., 1990 | 0.18 | 8.0 | 0.30 | 0.28 | - | - |

### C Perturbation analysis of three-state crossbridge model

We expand here the generalized two-state analysis to the three-state model by Murase et al., 1986 to identify the dominant time-constants in the system and compare them with the fitted values of Figure 5 - Table Supplement 1. In the three-state model, there are two attached states with bond distributions *n*_1_(*x, t*) and *n*_2_(*x, t*) and a detached state with distribution *n*_0_(*x, t*). By defining a column vector 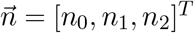, the governing dynamics of the system is given by

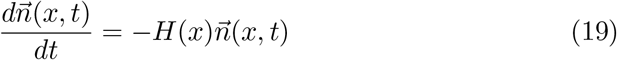

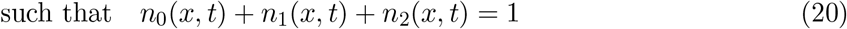

**Figure 3 - Figure Supplement 1:**
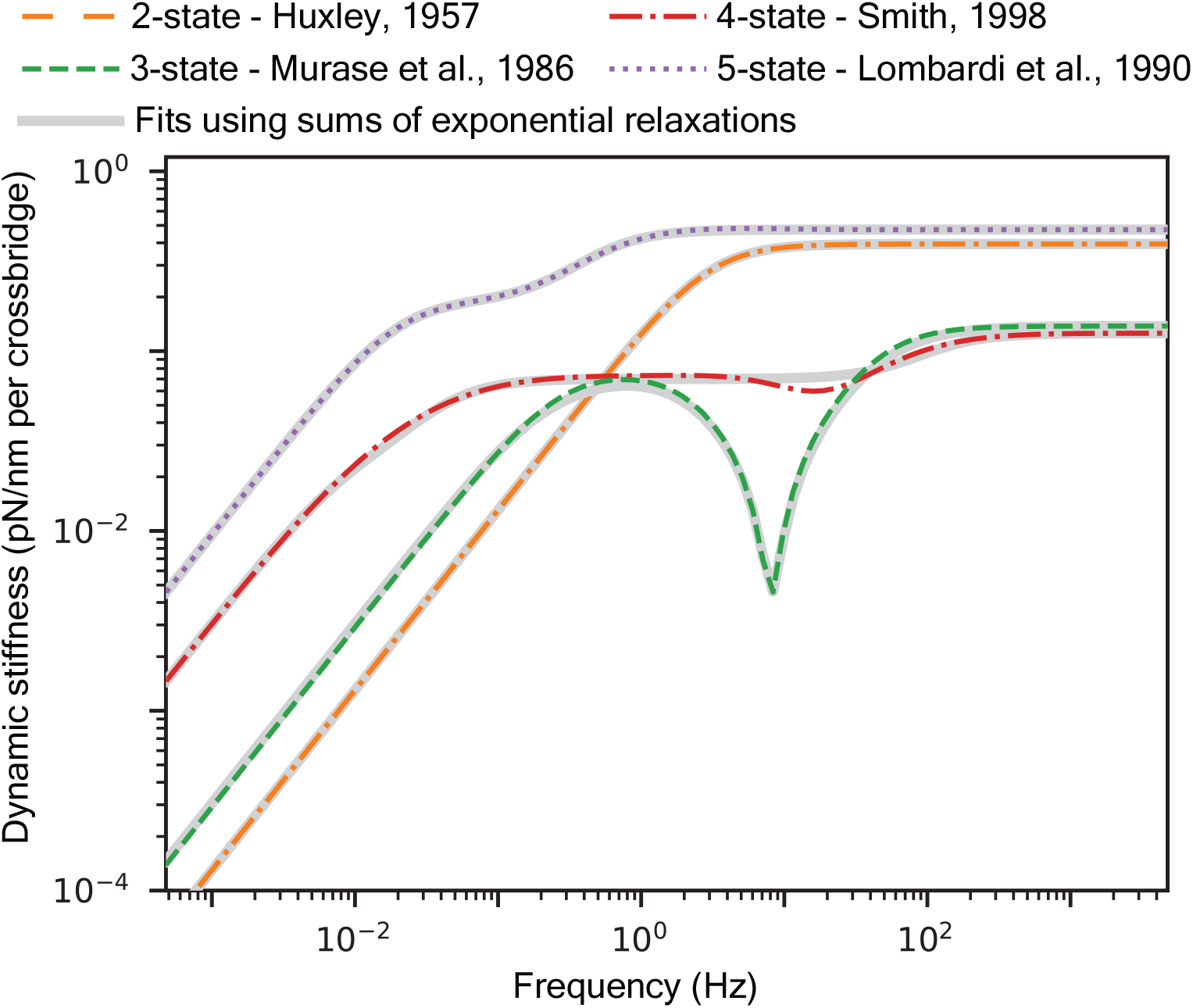
Fitted curves to the dynamic stiffness of different crossbridge models.

**Table Figure 5 - Table Supplement 2:** Comparison of fitted time-constants and time-constants identified by using local peaks in the derivatives of steady-state bond distributions for a two-state and a three-state crossbridge model. The peak of *xdn*_ss_/*dx* for the two-state model (equation (11)) is computed exactly with a time-constant *τ* = 1/(*f*_1_*h* + *g*_1_*h*) = 5.3*e* −2 seconds.

| Model | Fitted time-constants | Search using $x dn_{ss}(x)/dx$ |
| --- | --- | --- |
| Huxley, 1957 | 5.3e-2 seconds | 5.3e-2 seconds |
| Murase et al., 1986 | (0.59 , 6.8e-2 , 3.2e-3) seconds | (0.64 , 7.2e-2 , 3.7e-3) seconds |

and where *H*(*x*) is a 3×3 state transition matrix defined by the kinetic scheme of figure 5b. These equations uniquely define the steady-state bond distribution at equilibrium which we de-note as 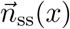. Specifically, the state transition matrix has three eigenvector-eigenvalue pairs 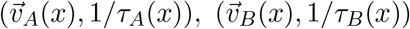, and 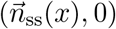 where *τ*_*A*_ and *τ*_*B*_ are the two time-constants driving the perturbation response and 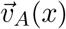 and 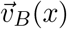 are unit vectors.

Mirroring the generalized two-state derivation, the perturbation response to a step length of size *a* is obtained by setting 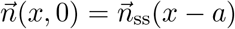 as the initial condition to arrive at

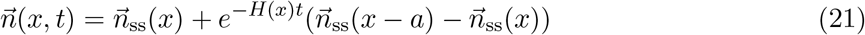

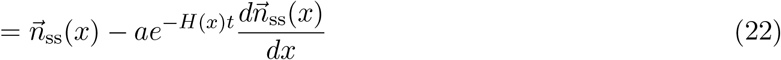

where the second equality arises by restricting our attention to small perturbations. We now expand the matrix-vector multiplication on the RHS using eigendecomposition of *H*(*x*) as

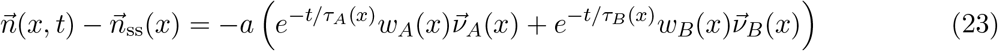

where *w*_*A*_ and *w*_*B*_ are linear weights that satisfy the linear system

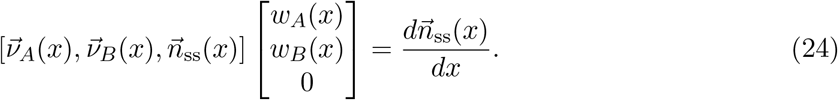

The third linear weight is necessary zero because 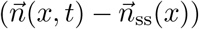 in equation (23) must decay to zero and cannot have a component parallel to *n*_ss_(*x*) which does not decay.

The perturbation response is the first moment of equation (23) multiplied by a stiffness vector that maps the bond distributions to forces. It is given in terms of crossbridge density *M*, stiffness of the first attached state *λ*_1_, and stiffness of the second attached state *λ*_2_ as

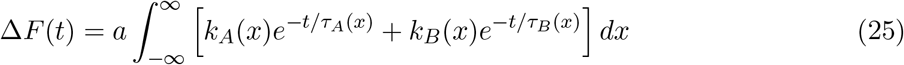

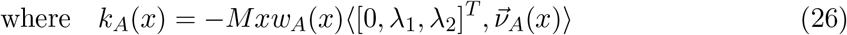

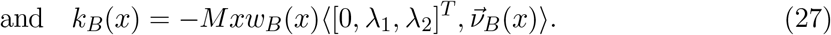

The operator ⟨·⟩denotes the dot product between two column vectors. The first entry in the stiffness vector is set to zero to represent the detached crossbridge state. Our analysis of the generalized two-state crossbridge model suggests that dominant time-constants appears where either *k*_*A*_(*x*) or *k*_*B*_(*x*) are localized, and we find that these time-constants do not significantly differ from heuristically using the derivatives of the steady-state distributions (figure 5 - Figure Supplement 1).

**Figure 5 - Figure Supplement 1:**
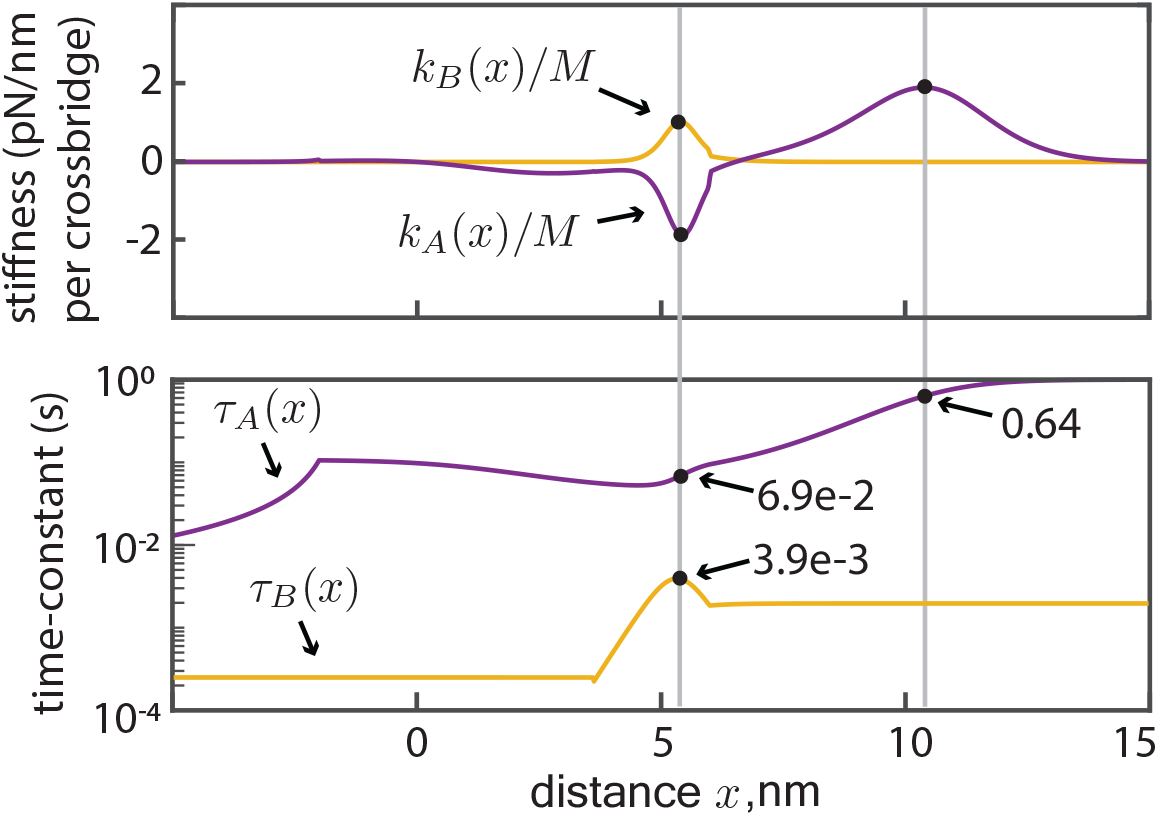
Examination of the local peaks in stiffnesses *k*_*A*_(*x*) and *k*_*B*_(*x*) identifies the dominant time-constants. The stiffnesses are computed using an eigenvector analysis of the state-transition matrix *H*(*x*) and are different from *λ*_1_*xdn*_1,ss_(*x*)/*dx* and *λ*_2_*xdn*_2,ss_(*x*)/*dx* used in figure 5c as a heuristic search. The identified time-constants do not significantly differ from the values found in figure 5c.

